# Comment on ‘Optimal Policy for Multi-Alternative Decisions’

**DOI:** 10.1101/2019.12.18.880872

**Authors:** James A. R. Marshall

**Affiliations:** University of Sheffield, Department of Computer Science

## Abstract

Optimality analysis has recently been proposed for value-based decision-making, in which decision agents are rewarded by the value of the selected option. This contrasts with psychophysics where decision agents are typically rewarded only if they choose the ‘correct’ or best option. The analysis of optimal policies for value-based decisions raises interesting and surprising parallels with decision rules proposed for accuracy-based decisions in binary and multi-alternative cases, and explains experimentally-observed deviations from rationality. However, the analysis assumes that decision agents should treat time as a linear cost, and thus optimise their Bayes Risk from decisions. A more naturalistic assumption is that future rewards are geometrically discounted, since they are less likely to be realised in an uncertain world. Changing the way in which time is costed leads to substantive changes in the resulting optimal policies, explains empirical data that previously could not be explained, and makes falsifiable predictions for future experiments.

## Introduction

In understanding the brain, which is a product of evolution, searching for optimal algorithms for typical decision problems can provide great insight. This normative approach can explain observed behaviours, and predict new behavioural patterns, based on evolutionary advantage. Yet the assumptions underlying such model analyses can prove crucial. In a recent article Tajima and colleagues ask what optimal decision algorithms look like for multi-alternative value-based choices, in which subjects are rewarded not by whether their decision was correct or not, but by the value to them of the selected option [Tajima et al., 2019]. The resulting algorithms correspond to earlier simple models for perceptual and value-based decision-making. Tajima *et al.*’s findings, however, rest on an assumption that time is a linear cost for subjects. When, as is more appropriate for naturalistic decisions, subjects discount rewards more the further into the future they will occur, optimal decision algorithms change qualitatively. These changes are consistent with recent empirical data that cannot be explained by analysis based solely on a linear cost of time.

In analysing multi-alternative value-based decision-making, Tajima *et al.* build on their earlier work in optimal decision policies for binary value-based choices [Tajima et al., 2016]. Through sophisticated analysis based on the standard tool for solving such decision problems, stochastic dynamic programming [Mangel and Clark, 1988, Houston and McNamara, 1999], the authors also seek neurally-plausible decision mechanisms that may implement or approximate the optimal decision policies [Tajima et al., 2016, Tajima et al., 2019]. These policies turn out at their simplest to be described by rather simple and well-known decision mechanisms, such as drift-diffusion models with decision thresholds that collapse over time for the binary case [Tajima et al., 2016], and nonlinear time-varying thresholds that interpolate between best-vs-average and best-vs-next in the multi-alternative case [Tajima et al., 2019].

## Results

The purpose of this commentary is not to criticise the methods used by Tajima and colleagues, which are standard, or the analyses, which are elegant. Rather, the purpose is to question one of their central assumptions and draw attention to the changes in conclusions that may result when it is altered. Tajima *et al.* make an assumption that appears widespread in psychology and neuroscience, that decision makers should optimise their Bayes Risk from such decisions; this is equivalent to maximising the expected value of decisions in which there is a linear cost for the time spent deciding [Bogacz et al., 2006, Pirrone et al., 2014]. For a lab subject in a pre-defined and known experimental design this may appear appropriate, for example because there may be a fixed time duration within which a number of decision trials will occur. However, an alternative and standard formulation of the Bellman equation, the central equation in constructing a dynamic programme, accounts for the cost of time by discounting future rewards geometrically, so a reward one time step in the future is discounted by rate γ < 1, two time steps in the future by γ^2^, and so on (see Supplementary Data). This is a standard assumption in behavioural ecology [Mangel and Clark, 1988, Houston and McNamara, 1999], in which discounting the future means that future rewards are not guaranteed but are uncertain, due to factors such as interruption, consumption of a food item by a competitor, mortality, and so on. Thus discounting the future represents the inherent uncertainty that animals must make decisions under in their natural environments, in which their brains evolved. The appropriate discount is then the probability that future rewards are realised, hence geometric discounting is optimal since probabilities multiply. Indeed there is extensive evidence of such reward discounting in humans and other animals (*e.g*. [Sellitto et al., 2010], although this frequently suggests hyperbolic rather than geometric discounting, a fact that in itself merits an explanation based on optimality theory [McNamara and Houston, 2009]).

In the following I show sample optimal policies for single-trial decisions when the change is made from linear costing of time, or Bayes Risk, to geometric discounting of future reward. In the binary case decision boundaries become non-linear (Fig. 1A to B), and ‘zip’ together over time (see Supplementary Data). Tajima and colleagues observed similar dynamics for the case of non-linear, saturating, utility functions for the decision maker ([Tajima et al., 2016], Fig. 6d), yet under geometric discounting, non-linear decision thresholds are inevitable even for linear subjective utility. Note that geometric discounting of future rewards is similar to, but not the same as, non-linear utility. In the multi-alternative case, on the other hand, the picture is more nuanced; moving from linear costing of time to geometric discounting of future rewards changes complicated time-dependent non-linear decision thresholds (Fig. 1C, [Tajima et al., 2019] Fig. 7) into either simple linear ones that collapse over time for lower-value option sets (Fig. 1D), or nonlinear boundaries that evolve over time similarly to the Bayes Risk-optimising case for higher-value option sets (Supplementary Data). As Tajima *et al.* note, the simpler linear decision boundaries implement the ‘best-vs-average’ decision strategy, whereas the more complex boundaries interpolate between ‘best-vs-average’ and ‘best-vs-next’ decision strategies [Tajima et al., 2019].

**Figure 1.**
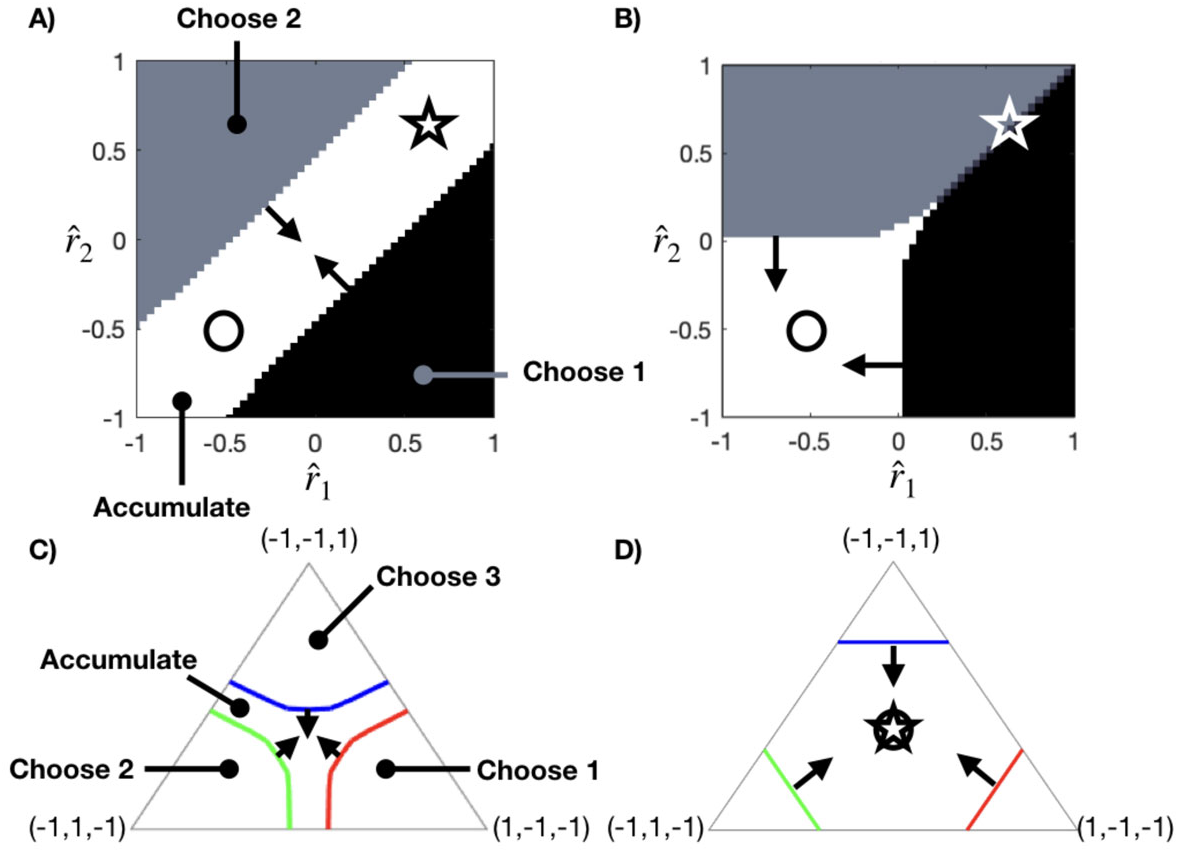
Optimal policies for value-based decision-making specifying when to choose options or continue accumulating evidence. A) For binary decisions with linear subjective utility, the optimal policy to maximise expected reward is a drift-diffusion model with time-collapsing boundaries [Tajima et al., 2016, Tajima et al., 2019]. This policy cannot explain observations of magnitude-sensitive reaction times, such as decisions between equal-but-low-value options (circle) being made faster than equal-but-high-value options (star) [Pirrone et al., 2018a]; this is because the decision-boundaries of the optimal policy have slope 1 in expected reward space 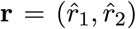, and true values for equal alternative decision pairs all lie on the line with slope 1, hence collapsing decision boundaries will intersect with both low-value and high-value pairs at the same time on average. B) In contrast, for binary linear utility decisions with geometric discounting of future rewards the optimal policy realises non-linear decision boundaries that ‘zip’ together (see Materials and Methods and Supplementary Data); such a policy can explain observed reaction time patterns as equal-but-high-value decision options (star) will be spontaneously chosen between. C) For multialternative decisions with linear utility, the optimal Bayes Risk-optimising policy exhibits time-dependent decision boundaries in the estimate space 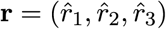; over time these boundaries interpolate between best-versus-average and best-versus-next decision strategies [Tajima et al., 2019]. D) In contrast, for multi-alternative decisions with linear utility, the optimal policy that maximises geometrically-discounted future rewards can exhibit simple linear decision boundaries that collapse over time for low-value option sets, corresponding to the best-versus-average decision strategy, or exhibit decision boundary dynamics more similar to the Bayes Risk-optimising strategy, in the case of high-value option sets (see Supplementary Data). As in the binary case, unlike the Bayes Risk-optimising strategy, maximisation of geometrically-discounted rewards predicts differing reaction times for equal-but-low-value decision option sets (circle), and equal-but-high-value decision options (star) due to faster collapse of decision boundaries in the latter case (see Supplementary Data).

Changing the optimal policy has consequences for optimality explanations of observed behaviour. For example, magnitude-sensitive reaction times have been observed in perceptual decisions by humans [Teodorescu et al., 2016, Pirrone et al., 2018a], and economic decisions by monkeys [Pirrone et al., 2018a], a phenomenon that has even been observed in single-celled organisms [Dussutour et al., 2019]. Pirrone *et al.,* for example, observed magnitude-sensitive reaction times when subjects were faced with pairs of equal options. This is incompatible with the ‘optimal’ policy assuming linear utility and Bayes Risk optimisation (Fig. 1A); under Tajima and colleagues’ analysis the explanations for such a behavioural pattern are either non-linear subjective utility, or learning about option value distributions over repeated trials [Tajima et al., 2016]. The latter can be discounted as single trial experiments also exhibit magnitude-sensitivity [Pirrone et al., 2018b], leaving saturating utility and discounting of the future as the principal remaining explanations. Distinguishing these empirically may be hard since singly or jointly they give rise to qualitatively very similar predictions (see Supplementary Data).

## Discussion

### Behavioural Predictions

The re-consideration of behavioural predictions for binary decisions highlights the need to re-evaluate predictions made by Tajima *et al.* for multialternative decisions, when Bayes Risk is replaced with discounting of future rewards; the Bayes Risk optimal policy is approximated by a neural model that is consistent with observations of economic irrationality [Tajima et al., 2019], hence it will be important to see if a revised neural model based on the revised optimal policy still shows such agreement. For example, while in the binary case magnitude-sensitive reaction times can be explained both by nonlinear subjective utility functions, and by geometric discounting rather than Bayes Risk, in the multi-alternative case preliminary exploration suggests that the same phenomenon is explained *only* by geometric discounting of future rewards (Fig. 1D) and not by nonlinear utility, at least for plausible nonlinear functions examined (see Supplementary Data). If generally true, magnitude-sensitive reaction times could thus falsify the Bayes Risk-optimisation account of behaviour, and further theoretical and empirical investigation into this would seem merited; if the analysis of [Tajima et al., 2019] showing magnitude-insensitive boundaries for linear utility and Bayes Risk-optimisation could be extended to determine conditions for utility functions to lead to magnitude-insensitivity, for example, this would be of great interest.

### Optimal Policies

The Drift-Diffusion Model optimises speed-accuracy trade-offs [Bogacz et al., 2006] yet has been criticised as being not generally applicable to value-based decisions [Pirrone et al., 2014]. Thus it is surprising when the optimal policy for value-based decisions is realised by a drift-diffusion process with collapsing thresholds [Tajima et al., 2016]. Here we have seen that with geometric discounting of future rewards the drift-diffusion model with collapsing bounds is not the optimal policy.

### Optimality Criteria

Practitioners of behavioural ecology have established principles to deal with empirically-observed deviations from the predictions of optimality theory [Parker and Smith, 1990]; two of the most useful are to consider that the optimisation criterion has been misidentified, or the behaviour in question is not really adaptive. Tajima and colleagues employ an exemplary approach, attempting to combine the best of the approaches of normative and mechanistic modelling [McNamara and Houston, 2009]; yet it bears remembering that subjects may not be trying optimally to solve the simple decision problem they are presented in the lab, but rather making use of mechanisms that evolved to solve the problem of living in their natural environment [Fawcett et al., 2014].

## Materials and Methods

Optimisation codes for the results presented here can be downloaded from https://github.com/DiODeProject/Optimal-policy-for-value-based-decision-making and https://github.com/DiODeProject/MultiAlternativeDecisions

An illustration of the optimal policy over time for binary decisions is given in Supplementary Data, as are illustrations of the effects claimed in the main text of option value on decision policies under geometric reward discounting with linear and nonlinear utility functions, and Bayes Risk optimisation with nonlinear utility functions.

## Acknolwedgements

I thank Satahiro Tajima for sharing the code for the binary decision model. Dr Tajima was an exceptionally promising scientist who is sadly missed. I thank Jan Drugowitsch and Alex Pouget for discussions over the content of this commentary. I thank Angelo Pirrone, Thomas Bose, Andreagiovanni Reina, Nathan Lepora and Sophie Baker for comments on an earlier draft.

## Supplementary Data

### 1. Bellman Equation for Maximisation of Geometrically-Discounted Reward

The revised Bellman equations (*cf.* Eq. 1, [Tajima et al., 2019]) used in the dynamic programme (implemented in the code found in the GitHub repositories cited in the main text) follow the general pattern

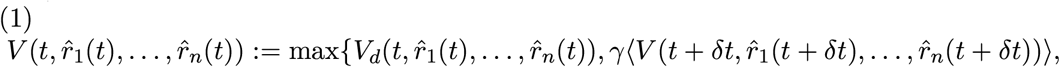

where *V* is the expected value at time *t* given estimator states 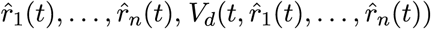 is the expected value of making a decision at time *t* given estimator states 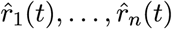 (defined as the value of the largest estimator), γ is a discount factor between 0 and 1, and (…) computes expectation marginalised over future estimator state probabilities.

### 2. Binary Decisions: Decision-Boundaries ‘Zip’ Together for Binary Decisions Under Geometric Discounting

The optimal time-dependent policy for geometric discounting is illustrated in the video https://github.com/DiODeProject/Optimal-policy-for-value-based-decision-making/blob/master/geometric.m4v.

### 3. Multi-Alternative Decisions: Optimal Policies are Magnitude-Sensitive Under Geometric Discounting

While it is proven that for Bayes Risk-optimisation decision boundaries are parallel to the diagonal passing through (0,0,0) and (1,1,1) [Tajima et al., 2019], for geometrically discounted rewards increasing the value of a set of equal options (which moves the relevant decision triangle along that diagonal as shown in Fig. 1) magnitude-sensitive reaction times manifest (in Fig. 2 decision boundaries collapse faster hence equal-alternative options in the centre of the triangle will be hit sooner by a decision boundary as their values increase).

### 4. Multi-Alternative Decisions: Optimal Policies Appear Magnitude-Insensitive for Selected Nonlinear Utility Under Bayes Risk-Optimisation

Under Bayes Risk-optimisation in trinary decisions it is known that optimal policies are magnitude-insensitive when subjective utility is linear [Tajima et al., 2019], whereas for binary decisions Bayes Risk-optimisation leads to magnitude-sensitivity when subjective utility is nonlinear [Tajima et al., 2016]. Under reasonable nonlinear subjective utility functions, however, optimal policies for Bayes Risk-optimisation over three options appear magnitude-insensitive (Fig. 3).

As can be seen from Fig. 3, the magnitude-insensitive strategy with utility defined according to a hyperbolic tangent function is simple ‘max’ with no accumulation of evidence. As can be also be seen from Fig. 3, the magnitude-insensitive strategy with utility defined according to a square root function is closer to the ‘max-vs-next’ strategy observed in [Tajima et al., 2019].

### 5. Multi-Alternative Decisions: Optimal Policies Become Magnitude-Sensitive Again for Nonlinear Utility Under Geometric Discounting

While section 4 showed that selected nonlinear utility functions, unlike the binary decision case, do not lead to magnitude-sensitivity in multi-alternative decisions with Bayes Risk-optimisation, here we show that simply switching from Bayes Risk to geometric discounting of future rewards reintroduces magnitude-sensitivity to the optimal policies for those same utility functions; this is illustrated in Fig. 4. For both nonlinear utility functions considered, geometric discounting of rewards results in optimal policies very similar to those for Bayes Risk-optimisation (Fig. 2).

---

**Figure 1.**
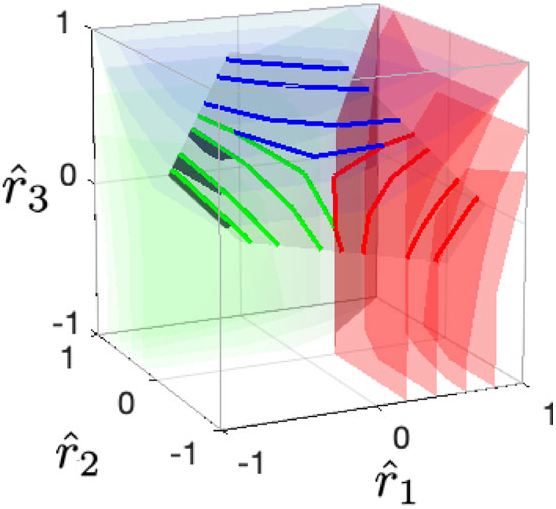
Geometric discounting of reward in trinary decisions - sample optimal policy with linear utility, discount rate γ = 0.8, value scale Δ*v* = 0.5.

**Figure 2.**
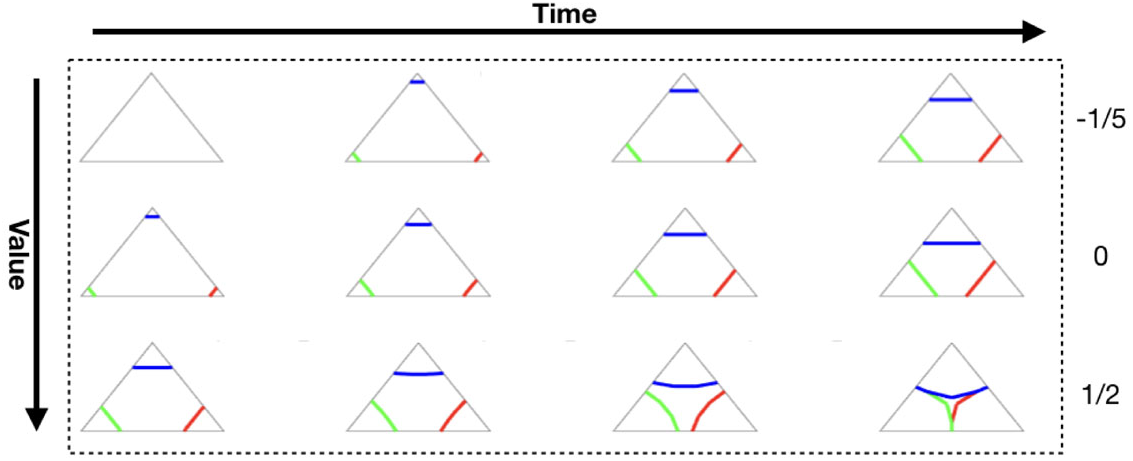
Geometric discounting of reward in trinary decisions - optimal policy with linear utility, discount rate γ = 0.8, value scales Δ*v* as indicated. Magnitude-sensitive reaction times are a feature of the optimal policy. For full explanation see Fig. 1 in the main text.

**Figure 3.**
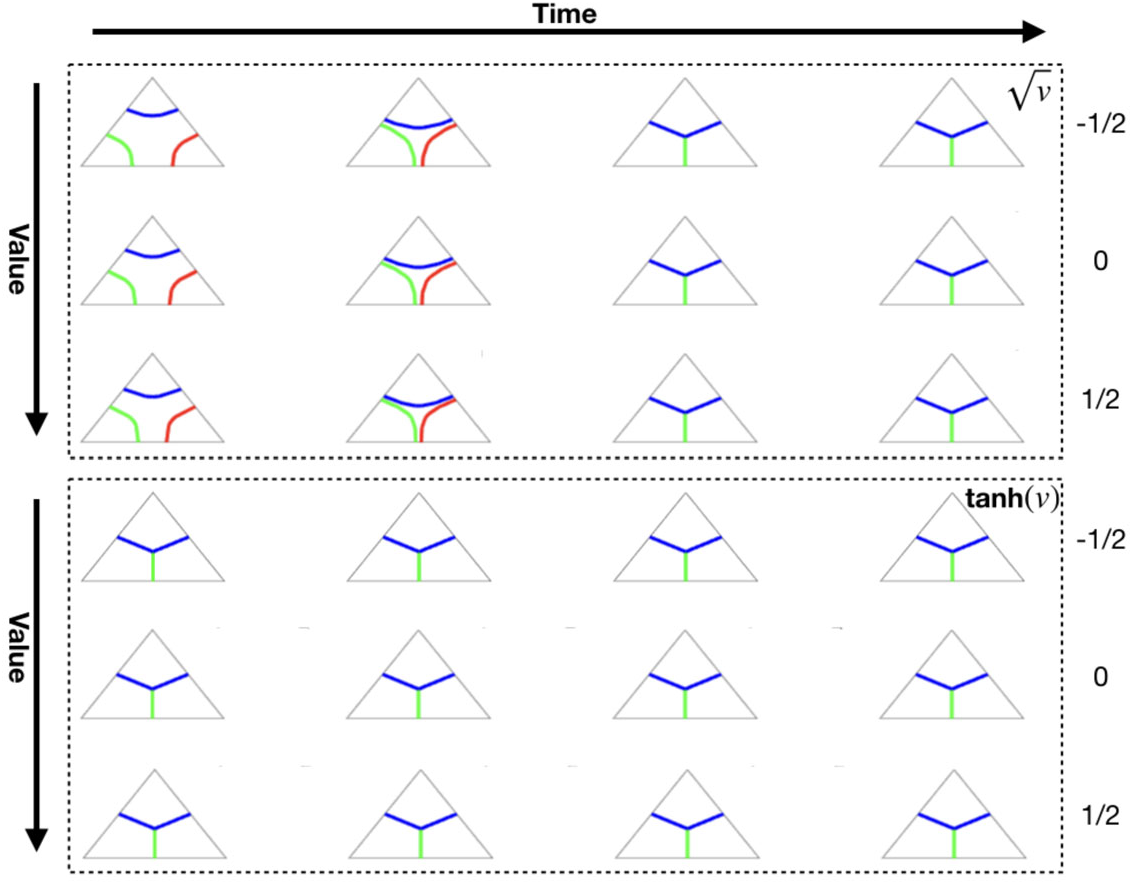
Bayes Risk in trinary decisions - optimal policy with nonlinear utilities 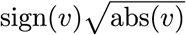 and tanh(*v*), value scales An as indicated. Utility functions indicated do not give rise to magnitude-sensitive reaction-times. For full explanation see Fig. 1 in the main text.

**Figure 4.**
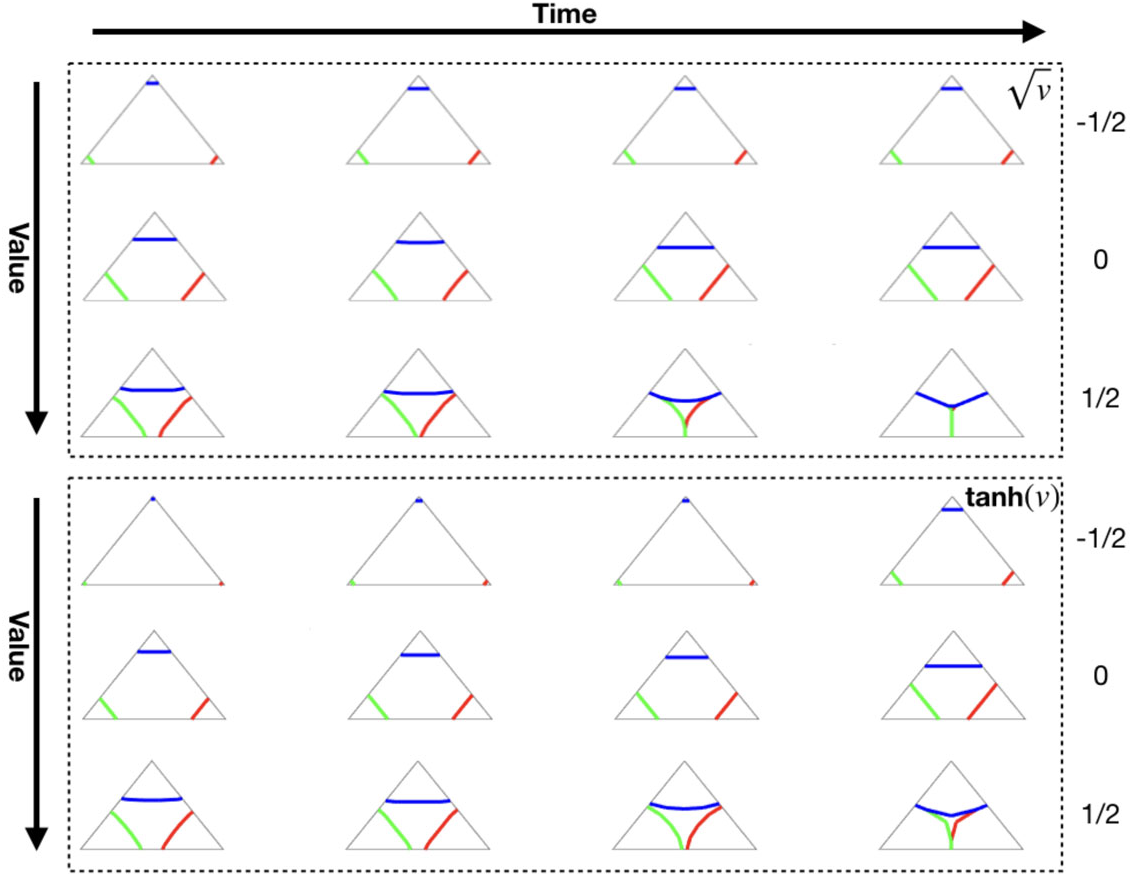
Geometric discounting of reward in trinary decisions - optimal policy with nonlinear utilities 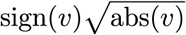 and tanh(*v*), value scales Δ*v* as indicated. Unlike the Bayes Risk case, optimal policies give rise to magnitude-sensitive reaction times. For full explanation see Fig. 1 in the main text.

## Notes

https://github.com/DiODeProject/Optimal-policy-for-value-based-decision-making

